# Alcohol-responsive genes identified in human iPSC-derived neural cultures

**DOI:** 10.1101/381673

**Authors:** Kevin P. Jensen, Richard Lieberman, Henry R. Kranzler, Joel Gelernter, Jonathan Covault

## Abstract

Alcohol use contributes to numerous diseases and injuries. The nervous system is affected by alcohol in diverse ways, though the molecular mechanisms of these effects are not clearly understood. Using human-induced pluripotent stem cells (iPSCs), we developed a neural cell culture model to identify the mechanisms of alcohol’s effects. iPSCs were generated from fibroblasts and differentiated into forebrain neural cells cultures that were treated with 50 mM alcohol or sham conditions (same media lacking alcohol) for 7 days. We analyzed gene expression using total RNA sequencing (RNA-seq) for 34 samples derived from 11 subjects and for 10 samples from 5 subjects in an independent experiment that had intermittent exposure to the same dose of alcohol. We also analyzed genetic effects on gene expression and conducted a weighted correlation network analysis. We found that differentiated neural cell cultures have the capacity to recapitulate gene regulatory effects previously observed in specific primary neural tissues and identified 226 genes that were differentially expressed (FDR< 0.1) after alcohol treatment. The effects on expression included decreases in *INSIG1* and *LDLR,* two genes involved in cholesterol homeostasis. We also identified a module of 58 co-expressed genes that were uniformly decreased following alcohol exposure. The majority of these effects were supported in the independent alcohol exposure experiment. Enrichment analysis linked the alcohol responsive genes to cell cycle, notch signaling, and cholesterol biosynthesis pathways, which are disrupted in several neurological disorders. Our findings suggest that there is convergence between these disorders and the effects of alcohol exposure.

## INTRODUCTION

Alcohol is commonly consumed worldwide^1^. Although moderate intake of alcohol may have modest health benefits^2, 3^, its misuse significantly contributes to numerous diseases and injuries from accidents^1, 4^ Alcohol consumption can progress to the development of an alcohol use disorder (AUD). AUD affects nearly 14% of the U.S. population and is characterized by tolerance to alcohol’s effects, continued use despite adverse consequences, and the development of withdrawal symptoms upon reducing alcohol intake^5, 6^. Persistent heavy alcohol intake has deleterious effects on the brain including changes in connectivity that are associated with a decline in cognitive abilities^7^, frontal lobe grey matter volume loss^8^, loss of white matter and neurodegeneration^9^, increased risk of early-onset dementia^10^, and decreases in cognitive function even with modest alcohol consumption^11^, among others^9,12-14^ Research suggests that the frontal cortex is particularly vulnerable to the degenerative effects of alcohol with large neurons being primarily affected ^8,15^. This population of neurons is also vulnerable in Alzheimer’s disease and normal aging^16,17^. It remains to be fully elucidated how the structural changes in the brain may relate to the development and progression of AUD.

Our understanding of the effects of alcohol at the molecular level in human neural cells is limited, due in part to the challenges associated with obtaining and culturing neural cells from human donors to perform controlled experiments. Differentiation of neural cell cultures from induced pluripotent stem cells (iPSCs)^18^ may provide an *in vitro* model to examine the effects of alcohol on human neural cells derived from characterized donors, potentially facilitating the identification of novel pathways associated with the effects of alcohol exposure on the brain. Recent work has highlighted the potential of iPSC technologies to study the complex actions of alcohol (for review, see^19, 20^). To our knowledge, no report to date has coupled human iPSC neural differentiation with RNA sequencing to explore transcriptome-wide effects of alcohol exposure *in vitro.*

The goal of the current study was to use RNA-Seq to characterize the effects of alcohol in neural cell cultures derived from iPSCs, which were differentiated into forebrain-type neural cells enriched for glutamatergic neurons. Two cohorts were utilized in the current study; a primary cohort of 34 neural cell cultures (from 10 donor subjects) exposed to alcohol continuously for 7 days, and a secondary cohort consisting of 10 neural cultures (from 5 donor subjects) exposed to a 7-day intermittent exposure to alcohol protocol. Our experimental approach included differential expression analysis and weighted gene co-expression-analysis to identify genes and pathways affected by alcohol exposure. With the bulk of our findings being consistent across two independent experiments, our findings highlight a role for genes involved in cholesterol homeostasis, notch signaling and cell cycle.

## MATERIALS AND METHODS

### Human induced pluripotent stem cells (iPSCs)

Fibroblasts obtained from subjects enrolled in studies at the University of Connecticut Health Center (Farmington, CT)^21-23^ were reprogrammed to iPSCs using CytoTune retro-or sendai virus kits (Thermo Fisher Scientific) by the University of Connecticut Stem Cell Core (Farmington, CT) and cultured on irradiated mouse embryonic fibroblasts as we have previously described in detail^24-26^. Fibroblast cultures tested negative for mycoplasma contamination. Informed consent was obtained from all subjects, and the study was approved by the University of Connecticut Health Center Institutional Review Board. Pluripotency of the selected colonies was verified by positive immunocytochemistry staining for SSEA-3/4 and NANOG by the UCONN Stem Cell Core. The primary analysis was based on a sample of iPSCs derived from 10 donor subjects (5 control and 5 AUD subjects). The primary sample included two clones that were selected from 1 AUD subject and one clone that was selected from each of the remaining 9 donor subjects, which yielded 11 independent iPSC lines. The second experiment was based on a set of iPSCs derived from 5 donor subjects (2 control and 3 AUD). One of the control donors in the second experiment was also used in the primary experiment. Combined, our sample sets included iPSCs generated from 6 controls and 8 AUD donors. The average age of the control donors was 35 years and of the AUD donors 46 years. All donor subjects were male. A matrix describing the sample donors, sample preparation and analyses is shown in **Table S1**.

### Neural differentiation and Immunocytochemistry

iPSCs were differentiated into neural cell cultures utilizing an embryoid-body-based protocol that we have previously described in detail^24^. In the absence of specific morphogens, the protocol yields forebrain-type neural cell cultures enriched for glutamatergic neurons^27^. Following differentiation and plating onto matrigel-coated glass coverslips, neural cells were cultured and matured for 12 weeks prior to experimentation. Our prior work has demonstrated that 8-12 weeks of growth under this protocol generates neural cultures with functional electrophysiological properties as evidenced by mature action potentials, spontaneous synaptic activity, and expression of ligand-gated ionotropic receptors^24, 25^. Neural cell markers were examined in differentiated iPSC lines 12 weeks after plating via immunostaining, as we have described.^25^ Cells were fixed in 4% paraformaldehyde, permeabilized using 0.2 % triton X-100 (Sigma-Aldrich), and blocked in 5% donkey serum (Jackson ImmunoResearch). The following primary antibodies were diluted in 5% donkey serum and incubated for 24-48 hours at 4° Celsius: mouse anti-beta III-tubulin (1:500, MAB1637, Millipore), mouse anti-GFAP (1:500, MAB360, Millipore), and rabbit anti-MAP2 (1:500, AB5622, Millipore). Appropriate donkey anti-mouse alexa fluor 594 (1:1000, Life Technologies) and donkey anti-rabbit alexa fluor 488 (1:1000, Life Technologies) secondary antibodies diluted in 3% donkey serum were used prior to mounting in DAPI-containing media for visualization.

### Alcohol treatment

Following 12 weeks of maturation, media was fully replaced with either normal neural differentiation media (henceforth referred to as the sham condition) or media supplemented with 50 mM ethanol. Two experimental protocols were used. In the primary experiment, alcohol-containing or sham neural differentiation media was fully replaced every 24 hours (our prior work demonstrated that alcohol concentrations decrease from 50 mM to 18 mM after 24 hours of incubation).^24^ In the second experiment, media was fully replaced every 48 hours. In both experiments, neural cells were treated with alcohol-containing or sham media for 7 days. The primary cohort consisted of 17 iPSC lines differentiated and exposed to sham or alcohol. This includes iPSC lines derived from 3 control and 3 AUD donors that were differentiated into neural cultures on two separate occasions and exposed to sham or alcohol, and 2 control and 3 AUD lines differentiated once and exposed to sham or alcohol. The second experiment included of 5 iPSC lines (2 control and 3 AUD) differentiated once and exposed to sham or alcohol.

### RNA sequencing

After alcohol or sham treatment, RNA was extracted using TRIzol Reagent (Thermo Fisher Scientific) per the manufacturer’s protocol. The primary analysis was based on RNA pooled from 136 neural cell cultures, with RNA from 4 wells of a 24-well plate per condition (sham and alcohol-treated) per subject, pooled as input to generate 34 cDNA libraries (17 sham-treated, 17 alcohol-treated) for sequencing. The primary analysis included 24 samples that were ribosomal RNA depleted and 10 samples that were poly(A) enriched (**Table S1**). We refer to the ribosomal RNA-depleted samples as Batch 1 and the poly(A) enriched samples as Batch 2. The second experiment included RNA from 60 neural cell cultures, with 6 wells of a 24-well plate per condition (sham and alcohol treated), per subject, pooled as input to generate 10 cDNA libraries for sequencing. All of the samples in the second experiment were ribosomal RNA depleted. All RNA samples were treated with DNase I (Thermo Fisher Scientific). RNA Integrity Numbers were assessed prior to library preparation, and they ranged from 6.6 to 9.7 (mean= 8.5) for experiment 1 (Batch1 + Batch2) and ranged from 6.3-10 (mean =8.97) for experiment 2. Randomly primed cDNA libraries (200-to 500-bp inserts) were prepared and sequenced at the Genomics Core of the Yale Stem Cell Center using Illumina TruSeq chemistry for library preparation and the Illumina HiSeq 2000 platform to generate 100-bp reads. Samples from Batch 1 and from the second experiment were paired end sequenced, while Batch 2 was not. Batch 1 and 2 were each separately aligned to the hg19 version of the human reference genome using TopHat2^28^. The second experiment was aligned to the hg38 genome build using STAR^29^. To increase mapping uniformity among samples in the second experiment, cutadapt was used to remove Illumina adapters from the sequence reads prior to alignment^30^. RNA sequencing data are available via the Sequence Read Archive (SRA accession numbers: SRP154768, SRP154763, SRP154762).

### Variant calling and eQTL analysis

Genetic variants were called from the aligned RNASeq data using a Genome Analysis Tool Kit (GATK) workflow that included ‘SplitNCigarReads’ to improve mapping near exon–intron junctions and base recalibration^31^. Variants were called using GATK HaplotypeCaller for 1000 Genomes Phase 3 variants with hg19 reference genome sequence^31, 32^. The variant calling was restricted to reads that mapped uniquely to the genome. The called variants were filtered to exclude multi-allelic variants, variants with read depth < 30 and fisher strand values < 30.0, and clusters of 3 or more SNPs within a window of 35 bases. PLINK was used to convert VCF files to binary format with a filter to exclude variants with genotype quality (gq) score < 30^33^. For each sample pair (sham and alcohol), we used the genetic data from the sample with the highest call rate for eQTL analysis. For eQTL analysis, we focused on Batch 1 samples (ribo-depleted) because Batch 2 (poly(A) enriched) had a small effective sample size (2 unique samples). Variants with minor allele frequency < 0.2 and missing in > 50 % of the sample were excluded from the eQTL analysis. A principal component analysis examining the SNP variation among the samples showed tight clustering of samples from the same subject relative to other samples (**Figure S1**). The association of SNPs to gene expression was tested using Matrix eQTL^34^. We focused on identifying “cis” acting eQTLs that were < 1000 base pairs from the start and end of a gene based on the gene boundaries defined by the UCSC Genes track of the UCSC genome browser (https://genome.ucsc.edu). Genes that had greater than 2 counts per million reads in at least 13 Batch 1 samples (> 50% of total) were retained for eQTL analyses. Two CPM was roughly 12 counts for the sample with the smallest library size (6.0M) and 48 counts for the average (24.0M) library size. Included in the analysis model were treatment condition (sham or alcohol) and a variable representing the paired cultures (1 through 12). SNP effects on gene expression were tested with an additive linear model. A separate model was run to test for SNP by alcohol treatment interactive effects on gene expression. We tested 10,616 SNP–gene pairs for associations.

### Differential expression analysis

To characterize effects on gene expression, we analyzed the number of read counts per gene. For the primary analysis, read counts were determined using HTSeq with uniquely mapped reads and UCSC gene boundary definitions^35^. We used *voom* to test for differential gene expression based on alcohol versus sham treatment and AUD case versus control donor status^36^. Genes that had greater than 2 counts per million (CPM) reads in at least 18 samples (>50% of total) were retained for the analysis. Two CPM was roughly 12 counts for the sample with the smallest library size (6.0 million) and 48 counts for the average (24.0 million) library size resulting in a set of 13,258 transcripts. We hypothesized moderate effects on gene expression (fold change >1.5), which is consistent with our prior work on alcohol’s effects on gene expression in neural cultures derived from iPSC, and with 17 samples per treatment group we had > 80% power to detect effects at an FDR threshold of 10% ^37, 38^. For all analyses, counts were quantile normalized. For the alcohol treatment analysis, the duplicateCorrelation function of *voom* was used to account for correlations between paired cultures (sham and alcohol treated), while batch (ribo depleted (Batch1) or poly(A) enriched (Batch 2)) and treatment (sham or alcohol) were included as fixed effects. Case versus control effects were analyzed using the duplicateCorrelation function to account for correlations between repeated subjects with batch (ribo depleted (Batch1) or poly(A) enriched (Batch 2)), treatment (sham or alcohol), and case or control status included as fixed effects. A similar approach was used to analyze effects in the second experiment. For this analysis, gene counts reported by STAR were analyzed with an inclusion threshold of >3 CPM in > 5 samples, which was approximately 40 reads for the average library size (13.0 million), and a total of 12,233 transcripts. The duplicateCorrelation function of *voom* was used to account for correlations between paired samples and treatment (sham vs alcohol) was analyzed as a fixed effect.

### Weighted gene co-expression network analysis methods

We investigated the expression of groups of highly correlated genes using weighted gene co-expression network analysis (WGCNA)^39^. Prior to WCGNA network construction and module detection, Combat was used to adjust for differences between RNA-Seq Batch 1 (ribo-depleted) and Batch 2 (poly(A) enriched)^40^. The following parameters were used for WGCNA: network type=signed, soft power threshold=14, corFnc=“bicor”, maxP0utliers=0.05, minModuleSize=30, and modules were merged at maximum dissimilarity threshold of 0.25. The steps in the module construction are illustrated in **Figure S5**^39^. Eighteen modules were identified with sizes that ranged from 42 to 3263 genes (**Table S5**). The module eigengenes were analyzed for association to alcohol treatment using a mixed model that included treatment (sham or alcohol) as a fixed effect and a random effect to account for correlations between paired treatments (sham and alcohol). The threshold for significance was Bonferroni corrected (p < 0.003) to account for testing the association of 18 modules for effects of alcohol treatment.

### Pathway and enrichment analyses

Differentially expressed genes (adjusted p < 0.1) were evaluated with Ingenuity Pathway Analysis. Canonical Pathways with a Benjamini-Hochberg adjusted p value < 0.1 are reported. For this analysis, the genes were ranked by the log fold change reported by *voom.* We used the geneSetTest function of the Limma program to test whether gene sets identified in the primary sample were differentially ranked in the secondary sample. We tested for differential ranking of genes associated with alcohol treatment at an adjusted p value < 0.1, for 3 gene sets identified in the primary analysis, which were genes in the Notch Signaling pathway, Superpathway of Cholesterol Biosynthesis pathway and “yellowgreen” module and for sets of genes reported by a recent study that investigate the effect of alcohol on gene expression in rat brain^41^. We tested for differential ranking of eQTLs identified by GTEx among the eQTLs identified in the iPSC-derived neural cell cultures. For the iPSC-derived neural culture eQTLs results, if a SNP or gene was tested more than once, the SNP or gene with the lowest p value was retained for enrichment testing. This resulted in 4,834 non-redundant SNPs and genes. For tests of differential ranking we assumed a “mixed” effect direction, unless noted otherwise, and used the test statistic for ranking. We tested GTEx eQTLs from 10 brain regions, a non-neural tissue (whole blood), and a larger, composite list based on GTEx eQTLs identified in any tissue^42^. GTEx data were acquired from UCSC Genome Browser Tables (https://genome.ucsc.edu) based on the eQTLs from 44 Tissues from GTEx midpoint release (V6). We limited testing to SNPs that had the same allelic effect direction between iPSC and GTEx datasets. The overlap with GTEx summary data for each eQTL category is shown in **Table S7**.

## RESULTS

### iPSCs differentiate into frontal cortical-like neural cultures

We utilized an embryoid-body based differentiation protocol (**Figure 1A-F**) to generate mixed neural cultures from human iPSCs generated from control and AUD donors. Following 12 weeks of neural maturation, iPSC-derived cultures contained dense Beta III-tubulin positive neurites (**Figure 1G**), MAP2-positive neurons with pyramidal morphology (**Figure 1H-I**), and GFAP-positive astrocytes (**Figure 1I**). These findings are consistent with our prior work demonstrating this protocol efficiently generates mixed neural cell cultures with ~50% TBR1-positive glutamate neurons that have the ability to generate trains of action potentials, display spontaneous synaptic activity indicative of synapse formation, and express functional glutamatergic and GABAergic ionotropic receptors, with no difference in the ability of iPSCs from control or AUD donors to generate neural cultures^24, 25^.

**Figure 1.**
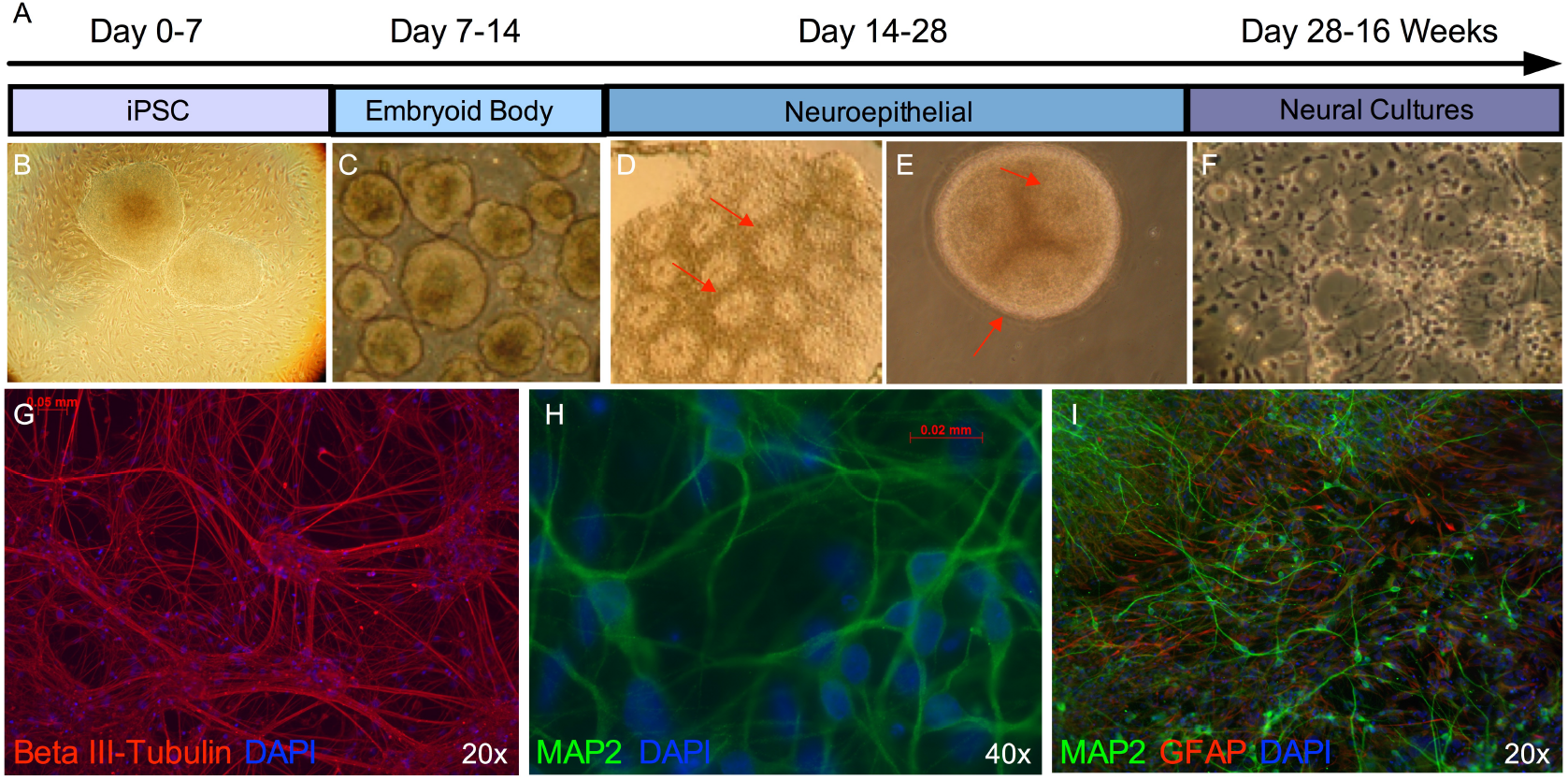
Neural differentiation of human iPSCs. (**A**) Schematic depicting the neural differentiation protocol. Induced pluripotent stem cells (**B**) are cultured on irradiated mouse embryonic fibroblasts for 7 days, following which they are cultured in suspension for 7 days to allow for the formation of embryoid bodies (**C**). Embryoid bodies are plated on a laminin substrate for 7 days to generate neuroepithelial cells (**D**), which form neural rosette-like structures (indicated by red arrows). Neuroepithelial cells are cultured in suspension for an additional 7 days to form and expand neurospheres (**E**), which display neural rosette-like structures (red arrows), before being manually dissociated and plated onto glass coverslips in neural media (**F**). After 12 weeks in neural media, cultures contain numerous Beta III-tubulin-positive neurites (**G**), pyramidal shaped MAP2-postive neurons (**H**), and GFAP-positive astrocytes (**I**).

To validate our neural differentiation, we used SNP information to identify gene regulatory effects in the neural cell cultures and characterized their relationship to effects previously reported for primary neural tissue. There were 14,770 autosomal exonic SNPs identified at a minor allele frequency greater than 20%, and that were genotyped in at least 50% of samples. In total, there were 10,690 SNP-gene association tests. There was 1 SNP-gene expression association significant at an FDR threshold of 5% and 72 SNP-gene associations significant at an FDR threshold of 10%. The top associations are shown in **Figure S2**. Notably, SNPs that were previously identified as eQTLs in cortex tissue had effects that ranked significantly higher among the eQTLs identified in iPSC-derived neural cultures (anterior cingulate cortex (BA24), p= 0.001; cortex, p=0.003; and frontal cortex(BA9)), p= 0.012; **Figure 2**). In contrast, SNPs that were previously identified as eQTLs from non-neural tissue (whole blood) were not ranked differently (p= 0.8). There was also little difference when examining a larger, composite list that included eQTLs previously identified in at least one of 44 tissues. The number of SNPs overlapping between datasets is show in **Table S7**. These findings indicate that the neural cell cultures have the capacity to recapitulate gene regulatory effects observed in primary tissue with some tissue specificity for frontal cortical regions.

**Figure 2.**
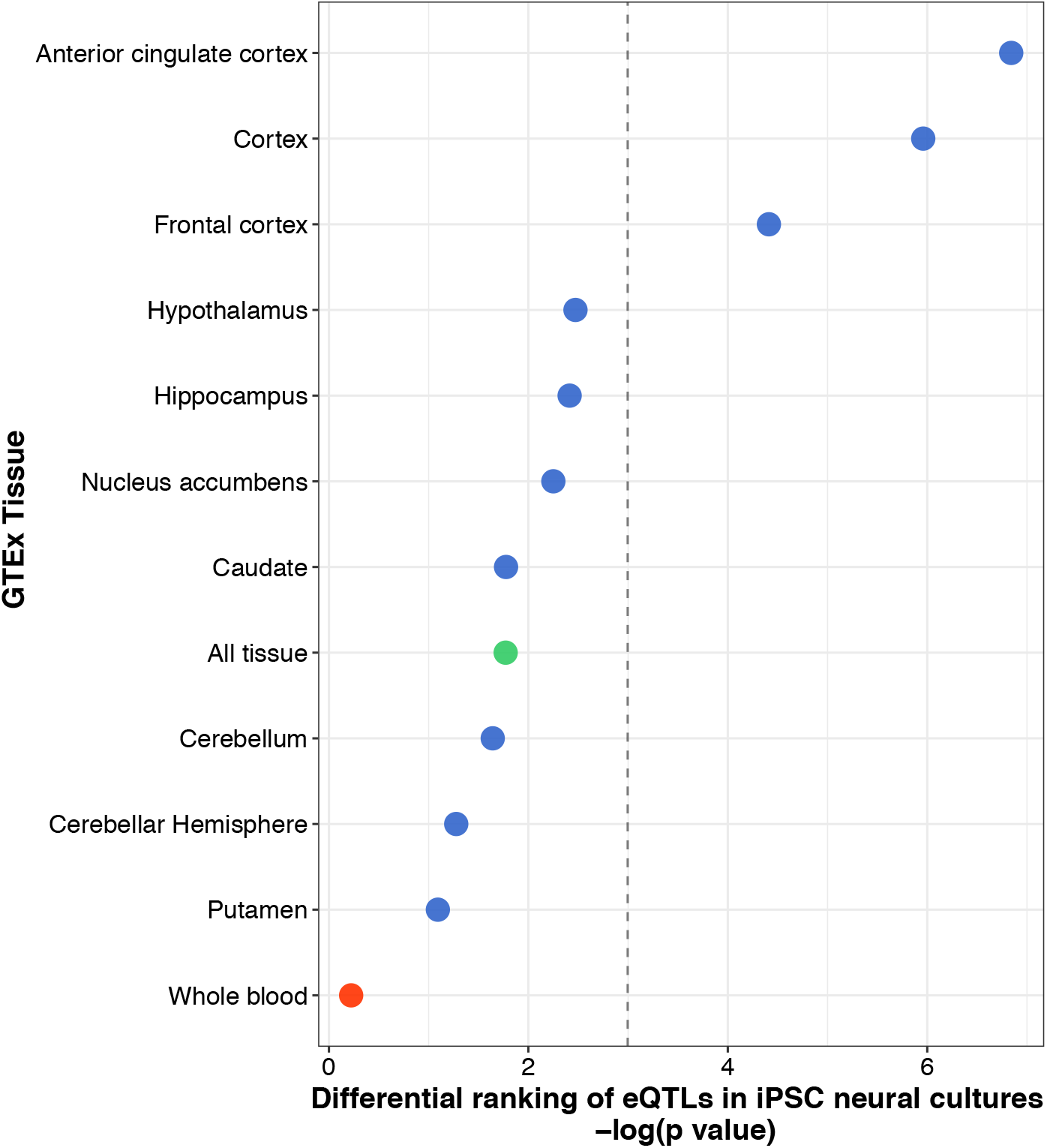
The effects of primary tissue eQTLs in neural cell cultures derived from iPSCs. The differential effects of previously identified eQTLs from 10 brain regions studied by the Genotype-Tissue Expression (GTEx) project. The effects in different brain regions (blue) are contrasted to a non-neuronal tissue, whole blood (red), and a composite set of previously identified eQTLs from 44 different tissues (green). The dashed line marks p < 0.05.

### The effects of alcohol on gene expression in neural cell cultures derived from iPSCs

We tested the effect of alcohol on gene expression following 7 days of alcohol or sham treatment. There were 13,258 transcripts that meet our inclusion criteria for testing. Alcohol treatment was associated with a change in the expression of 16 genes (adjusted p value < 0.05) and 226 genes at an adjusted p < 0.1. The gene with the top statistically ranked change was *INSIG1,* which encodes the Insulin-induced gene 1 protein. *LDLR,* which encodes the low-density lipoprotein receptor gene, was the second ranked change. The expression of both genes decreased in response to alcohol (**Table 1**). We examined the set of 226 genes (adjusted p < 0.1) in an independent alcohol treatment experiment that included 10 neural cell cultures with intermittent exposure to the same concentration of alcohol. In the second experiment, we observed consistent effects for genes that were either up or down regulated (p < 1×10^−5^) in the primary sample (**Figure 3**). We also used Ingenuity pathway analysis (IPA) to characterize the same set of 226 genes. The top IPA canonical pathway associations are shown in **Table S4**. The top ranked pathway was Notch Signaling, which included 5 genes. The effect directions for genes in **Table S4** are shown in **Table 1**. There were also several pathway associations related to cholesterol biosynthesis that were anchored on a common set of genes *(DHCR24, FDFT1, MSMO1).* Neither of the top two differentially expressed genes, *INSIG1* and *LDLR,* contributed to the associations to cholesterol biosynthesis pathways, despite what one might expect given their essential roles in cholesterol homeostasis^43^. As in the discovery sample, genes within the Notch Signaling pathway were uniformly decreased in the second experiment (p < 1×10^−4^), though genes in the cholesterol pathways and the Molybdenum Cofactor Biosynthesis pathway were not. There were no significant differences in gene expression between neural cell cultures derived from AUD and control subjects. There were also no significant SNP by treatment interactive effects (SNP x alcohol) associated with gene expression.

**Figure 3.**
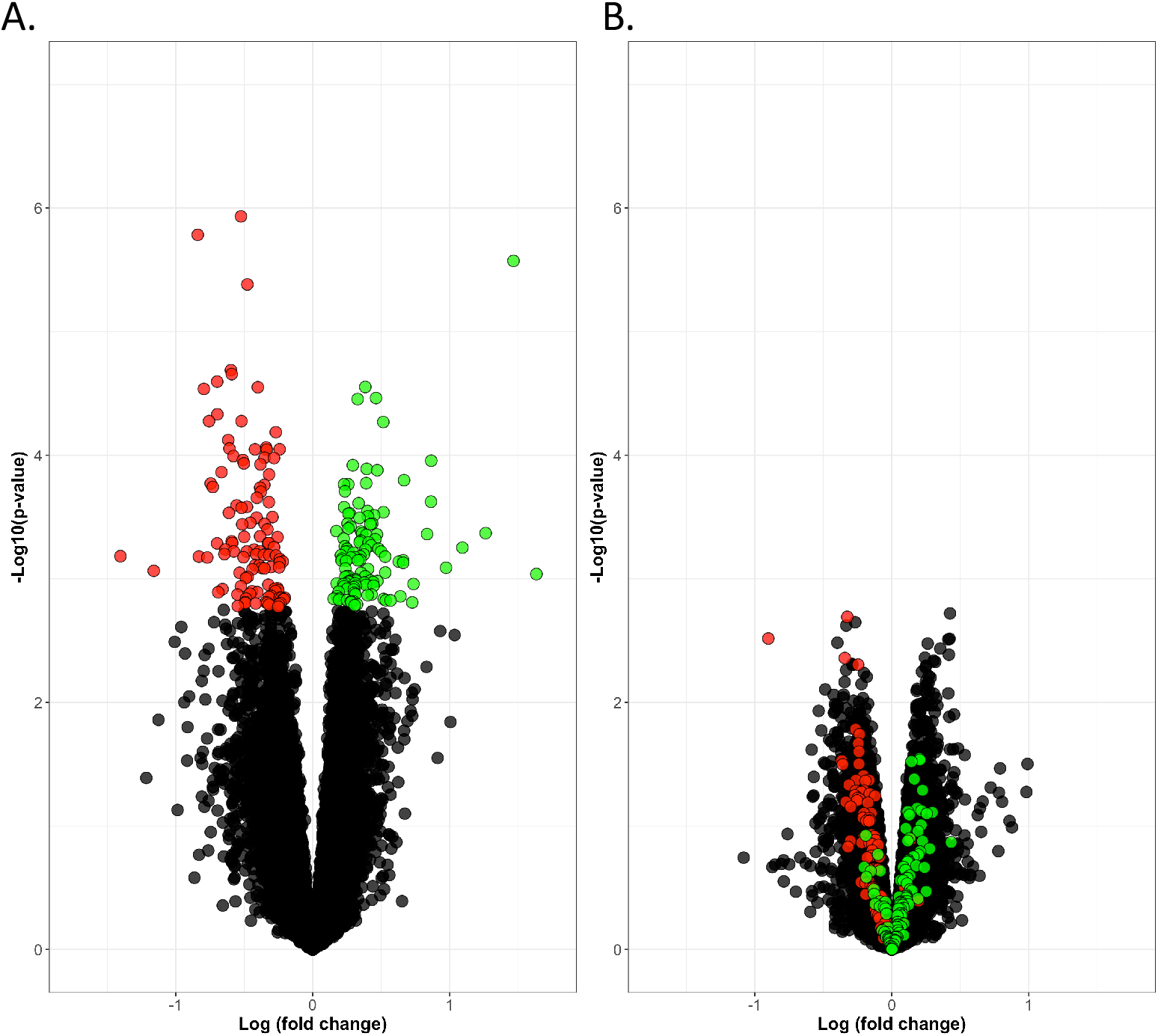
The effects of alcohol treatment on gene expression in neural cell cultures derived from iPSCs. There were 226 genes that were differentially expressed (p adj <0.1) in the discovery sample. The down (red) and up (green) regulated genes are show in A) the discovery sample (n=17 paired samples) and B) second sample (n=5 paired samples), which had intermittent exposure to the same concentration of alcohol.

**Table 1:**
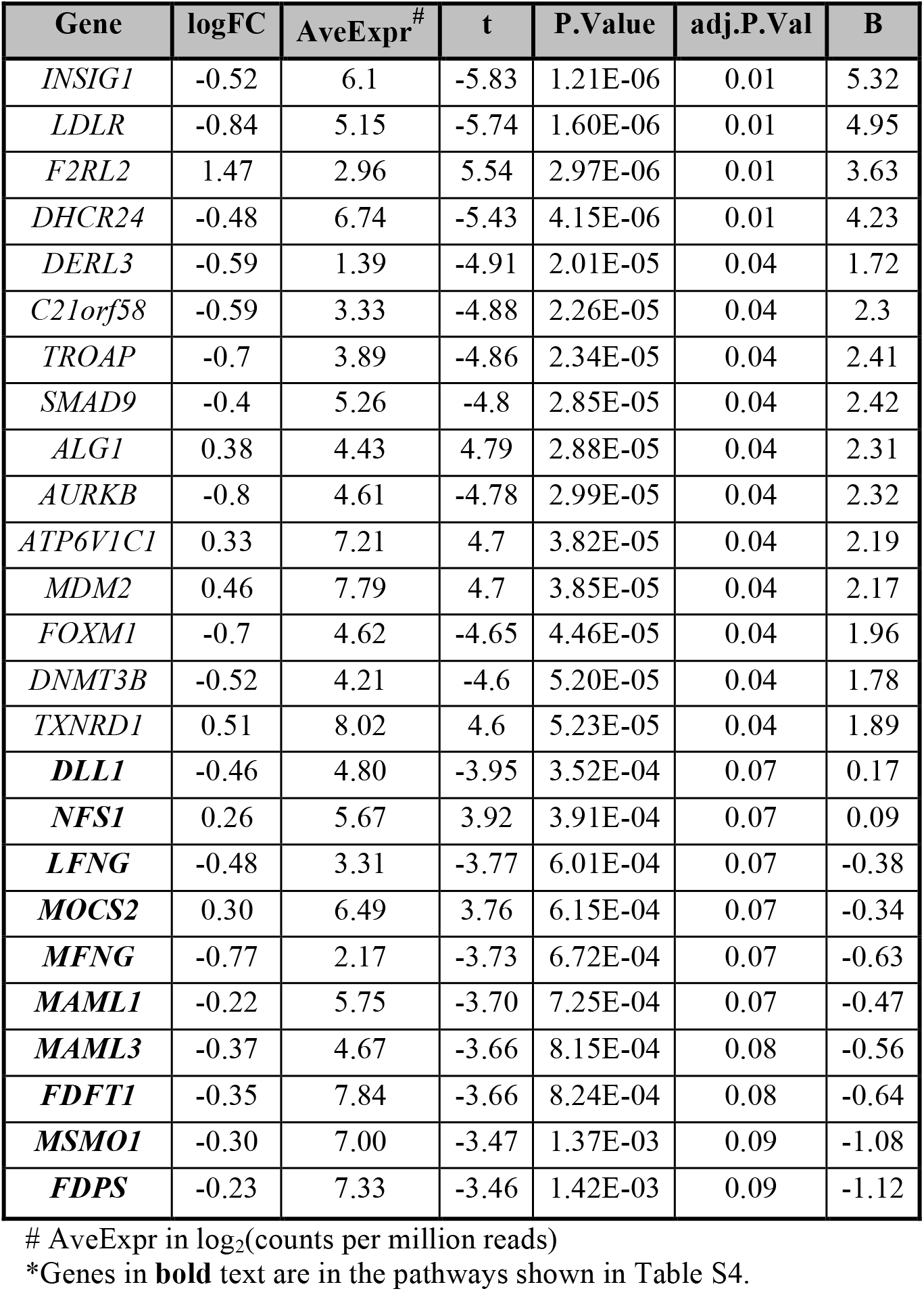
The effects of alcohol on differentially expressed genes with adjusted p < 0.05 and in Ingenuity canonical pathways.

Genes in the alcohol dehydrogenase gene family involved in the metabolism of ethanol, which are generally expressed at high levels in liver but not brain, where not highly expressed in the neural cell cultures, and 5 (out of 7) were excluded from the primary analysis because of very low expression levels. We conducted a secondary analysis of all mapped genes (no expression threshold), to characterize potential effects from this important gene family. The top effect in the primary and secondary experiment was for *ADH5.* There was a nominal increase in *ADH5* mRNA after alcohol exposure in the primary (unadjusted p < 0.05) and secondary (unadjusted p < 0.18) samples. *ADH5* was the highest expressed alcohol dehydrogenase gene in each sample, and it was included in the primary analysis. All alcohol dehydrogenase gene effects are shown in **Table S8**.

We also compared our results from the neural cell cultures derived from iPSCs to a recent study that investigated the effect of alcohol on gene expression in the ventral hippocampus and medial prefrontal cortex of adolescent alcohol-preferring rats^41^. Results (**Table S10** and **Figure S4**) showed that genes that were down regulated in ventral hippocampus were differentially expressed in the same direction in the alcohol-treated neural cell cultures derived from iPSCs (p=1.0×10^−6^). Genes that were up regulated in ventral hippocampus had a modest effect in the same direction (p= 6.8×10^−2^), whereas genes differentially expressed (up or down) in the rodent prefrontal cortex were not differentially expressed in iPSC. A group of 10 genes that had consistent evidence of up regulation in at least 4 out of 11 different rodent brain regions following alcohol exposure were also up regulated following alcohol treatment in the neural cell cultures (p= 2.2×10^−2^). Among these, the largest effect was for *ATF3* (log FC =0.43, p =0.012) followed by *BTG2* (log FC =0.21, p = 0.025). *DGKB,* the only gene that was down regulated in at least 4 out of 11 different rodent brain regions following alcohol exposure was unchanged in the neural cell cultures treated with alcohol.

### An alcohol responsive gene co-expression network

Using WGCNA, we identified eighteen modules that contained 42 to 3263 co-expressed genes. The module sizes are shown **Table S5**. The expression of one module, “yellowgreen,” was negatively correlated with the response to alcohol (Pearson r = -0.43, p = 0.0009), indicating that the expression of genes within this module was lower in the alcohol condition than the sham condition (**Figure 4B**). The 58 genes in the yellowgreen module were nearly uniformly decreased (p < 1×10^−6^) in the second experiment (**Figure 4C**). Within the yellowgreen module, the response to alcohol was correlated with the strength of the association to the module, i.e., genes that were more tightly associated with the module decreased more in the alcohol condition than genes that were less tightly associated to the module (Pearson r =-0.48, p= 0.00014) (**Figure S6**). Genes in the yellowgreen module are shown in **Table S6** DAVID analysis indicated that the yellowgreen module was strongly enriched with genes involved in the KEGG Pathway for Cell Cycle (FDR = 1.38×10^−4^) and for biological processes such as DNA replication, cell division, DNA repair, and DNA replication initiation (FDR < 3.44×10^−4^) (**Table S9**).

**Figure 4.**
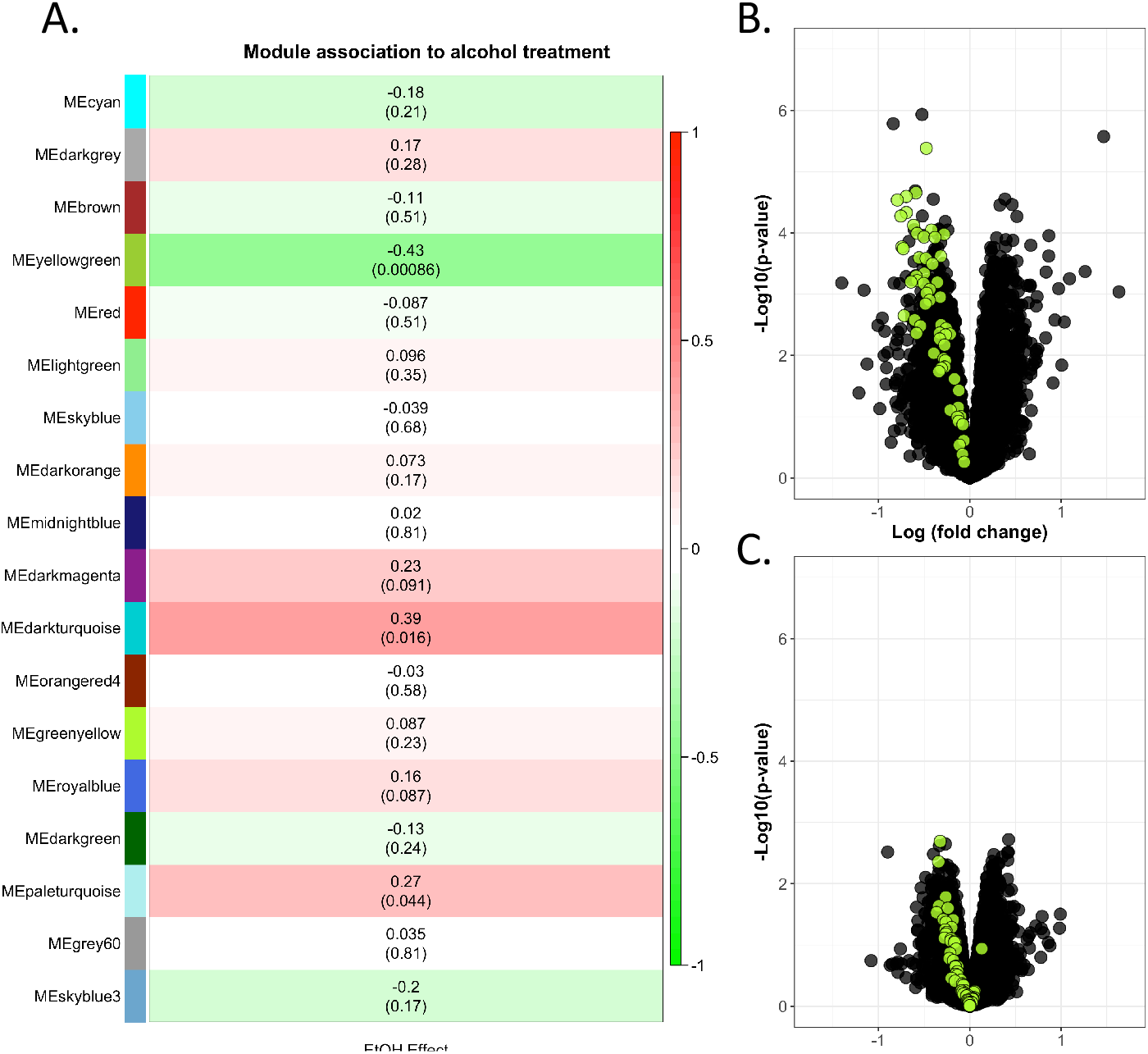
A module of co-expressed genes is down regulated by alcohol treatment. **A.** The expression of a module with 58 genes (“yellowgreen”) is lower in the alcohol treatment condition compared to sham. Shown are the Pearson correlations and p-values for the analysis of module expression to treatment condition. The color is weighted by the magnitude and direction of the Pearson correlation. The response to alcohol for genes in the yellowgreen module is shown in the primary experiment (**B**) and (**C**) second experiment.

## DISCUSSION

Our study used human neural cell cultures derived from iPSCs to characterize the effect of alcohol on gene expression. Our findings, the bulk of which were supported across two independent experiments, demonstrate that alcohol affects genes involved in cholesterol homeostasis, notch signaling, and cell cycle pathways. To complement our characterization of the neural cell cultures, we analyzed genetic effects on gene expression to demonstrate that the neural cell cultures have the capacity to recapitulate gene regulatory effects previously identified in primary neural tissues that are relevant to alcohol’s effects. Cholesterol homeostasis, notch signaling and cell cycle pathways are disrupted in several neurological disorders and our findings provide insight into the molecular basis for the potential convergence on these same pathways as a result of alcohol exposure.

Among the top differentially expressed genes following alcohol exposure were *INSIG1* and *LDLR,* which were both down regulated. Prior work has shown that alcohol exposure lowers LDLR levels in mice liver, where LDLR has a critical role in cholesterol turnover^44^. LDLR also has important functions in the brain. For example, LDLR overexpression reduces Aβ aggregation and neuro-inflammatory responses in a mouse model of Alzheimer’s^45^, and LDRL has also been reported to impact learning and memory^46^. Interestingly, *INSIG1* is also involved in regulating cholesterol in the cell, and genetic association studies suggests it has potential links to Alzheimer’s based a reported trend level genomewide association (p = 7×10^−7^) of an *INSIG1* 3’ untranslated region SNP to Alzheimer’s Disease in a family based association study^43, 47^. Along with changes for additional genes involved in cholesterol biosynthesis (**Table S4**), these findings indicate that alcohol may cause alterations to cholesterol homeostasis within the brain, which might be important clinically given research linking dysregulated cholesterol homeostasis to many neurological disorders, including Alzheimer’s Disease and dementia^48^. Also, cholesterol is a precursor to neuroactive steroids which are thought to mediate some of its behavioral effects via allosteric actions at GABA(A) receptor subtypes^49^. A recent clinical study demonstrated that inhibition of a key enzyme for neuroactive steroid biosynthesis reduces the subjective, sedative effects of acute alcohol intoxication^21^. Reduced sedation in response to alcohol exposure may be a risk factor for the development of AUD^50, 51^. Therefore, our finding that alcohol exposure perturbs the expression of genes regulating cholesterol homeostasis may help to explain the relationship between alcohol consumption, the development of AUD, and neurodegeneration.

Our analysis also indicates that several notch signaling pathway genes are affected by alcohol exposure. The notch signaling pathway, which is highly conserved among multicellular organisms, is active in the mammalian adult and developing nervous system. In the adult mammalian nervous system notch pathway genes have an important role in synaptic plasticity^52^, and in a drosophila model, mutations to genes within the notch signaling pathway disrupt the formation of memories for ethanol reward^53^. Given the important role of the notch signaling pathway in determining cell fate during development, our observations might relate to the detrimental effects of alcohol on the adult and/or developing nervous system^54, 55^. Likewise, control of the cell cycle and DNA replication, two pathways implicated by our WGCNA approach, are critical in the adult and developing nervous systems. Failure to maintain control of the cell cycle in adult neurons has been linked neurodegenerative disorders.^56^ Thus, these gene expression changes could relate to some of clinical observations that heavy alcohol intake is associated with neurodegeneration, a decline in cognitive abilities, and increased risk for early-onset dementia^7-9, 12, 13^.

Our findings are consistent with some prior studies of alcohol exposure that investigated effects in human and rodent primary neural tissue. For example, genes that were differentially expressed after alcohol exposure in the rat ventral hippocampus had similar effect directions in our human iPSC derived neural cell cultures treated with alcohol as a set of genes that had consistent effects in multiple different rat brain regions^41^. The McClintick et al. study^37^ also noted that alcohol exposure was associated with the down regulation of several cholesterol pathway genes, although not the same cholesterol pathway genes that we identified here. A study by Lewohl et al. that compared gene expression in post mortem frontal cortex from non-AUD and AUD subjects showed changes to some genes involved cell cycle regulation, results that are similar to our study’s^57^. These consistent effects are notable given prior studies that demonstrated limited overlap in differentially expressed genes from tissue collected at different developmental stages (e.g., adolescent vs adult) and from tissue from different brain regions at the same developmental stage. For example, in a study by Flatscher-Bader et al.^54^ comparing the nucleus accumbens and ventral tegmental area from AUD cases to controls, only 6% of the genes whose expression was associated with AUD were shared between the two tissues, and in a rodent study by McBride et al.^55^,there was limited overlap between differentially expressed genes from the nucleus accumbens shell and central nucleus of the amygdala of adolescent rats, and limited overlap when comparing the adolescent effects to effects previously identified in adults for the same tissues^58, 59^. The top alcohol dehydrogenase gene effect was *ADH5*. Although *ADH5* encodes a protein with limited alcohol metabolizing activity (K_m_ for ethanol > 1000), recent GWAS identified SNPs within *ADH5* associated with alcohol consumption, suggesting that *ADH5* has an important role in mediating the response to alcohol^60, 61^. It will be important in future studies to establish and characterize links between what is observed *in vitro* and in other models of alcohol exposure to the clinical manifestations of AUD.

Strengths of our study were the sample size (n=34) and the use of an independent sample (n=10), which was intermittently exposed to the same concentration of alcohol and could therefore reasonably be considered appropriate for replication, although the conditions were not identical. Our experimental design allowed for within-subject comparisons and our RNA-Seq samples were pools of multiple wells of a culture plate per condition and subject, which likely reduced potential sources of heterogeneity related to individual outliers. However, statistical power would still have been insufficient to detect certain effects; for example, we did not detect differences between neural cells from AUD cases and control subjects, nor did we detect eQTL effects that were modified by alcohol exposure. We elected to study the effects of 50 mM ethanol (equivalent blood alcohol concentration = 0.23 mg/dl), a concentration of alcohol that is commonly observed in individuals with moderate-to-severe AUD, but is higher than that resulting from low-to-moderate alcohol consumption. The cells in the second experiment had intermittent exposure to the same dose, and most of the effects on gene expression were consistent between the samples. Thus, the effects of exposure to 50 mM ethanol may to some degree generalize to lower levels of exposure, however additional studies on the effects of lower alcohol doses are warranted.

In conclusion, we used neural cell cultures derived from iPSCs to characterize the effects of alcohol on gene expression. We identified genes and pathways that are affected by exposure to alcohol, including cholesterol homeostasis, notch signaling and cell cycle. These effects point to molecular mechanisms that could contribute to alcohol-induced neurodegeneration. Clinically, alcohol-induced neurodegeneration can be profoundly debilitating, and it can complicate treatment efforts. With support from additional studies, the development of treatments that target these pathways could help to reduce the negative health effects associated with heavy alcohol use.

## Funding and conflicts of interest

Supported by NIH grants P60 AA03510 (Alcohol Research Center), AA23192 (HRK), AA015606 (JC), M01 RR06192 (University of Connecticut General Clinical Research Center), a US Department of Veteran Affairs Career Development Award, and by a grant from the CT Department of Public Health (JC).

Dr. Kranzler has served as a consultant, CME speaker, or advisory board member for the following companies: Indivior and Lundbeck. He is a member of the Alcohol Clinical Trials Group of the American Society of Clinical Psychopharmacology, which was supported in the last three years by Abbvie, Alkermes, Ethypharm, Indivior, Lilly, Lundbeck, Otsuka, Pfizer, Arbor Pharmaceuticals, and Amygdala Neurosciences. Drs. Kranzler and Gelernter are named as inventors on PCT patent application #15/878,640 entitled: “Genotype-guided dosing of opioid agonists,” filed January 24, 2018. All other authors declare no conflict of interest.

## Acknowledgements

We would like to thank Leann Crandall from the UCONN Stem Cell core for her valued assistance and expertise in generating iPSC lines.

